# Divergent gene expression patterns in alcohol and opioid use disorders lead to consistent alterations in functional networks within the Dorsolateral Prefrontal Cortex

**DOI:** 10.1101/2024.04.29.591734

**Authors:** Martha MacDonald, Pablo A. S. Fonseca, Kory Johnson, Erin M Murray, Rachel L Kember, Henry Kranzler, Dayne Mayfield, Daniel da Silva

## Abstract

Substance Use Disorders (SUDs) manifest as persistent drug-seeking behavior despite adverse consequences, with Alcohol Use Disorder (AUD) and Opioid Use Disorder (OUD) representing prevalent forms associated with significant mortality rates and economic burdens. The co-occurrence of AUD and OUD is common, necessitating a deeper comprehension of their intricate interactions. While the causal link between these disorders remains elusive, shared genetic factors are hypothesized. Leveraging public datasets, we employed genomic and transcriptomic analyses to explore conserved and distinct molecular pathways within the dorsolateral prefrontal cortex associated with AUD and OUD. Our findings unveil modest transcriptomic overlap at the gene level between the two disorders but substantial convergence on shared biological pathways. Notably, these pathways predominantly involve inflammatory processes, synaptic plasticity, and key intracellular signaling regulators. Integration of transcriptomic data with the latest genome-wide association studies (GWAS) for problematic alcohol use (PAU) and OUD not only corroborated our transcriptomic findings but also confirmed the limited shared heritability between the disorders. Overall, our study indicates that while alcohol and opioids induce diverse transcriptional alterations at the gene level, they converge on select biological pathways, offering promising avenues for novel therapeutic targets aimed at addressing both disorders simultaneously.

## Introduction

Substance use disorders (SUDs) are a chronic, relapsing diseases characterized by compulsive drug seeking despite severe harm or a desire to reduce or quit use [1]. Alcohol and Opioid Use Disorders (AUD/OUD) are two prevalent SUDs with the highest mortality rates among misused substances. AUD and OUD contribute to approximately 178,000 and 80,000 U.S. deaths annually, respectively [2, 3]. The annual economic toll of AUD and OUD is estimated to be US$720 billion, encompassing healthcare expenditures, lost productivity, collision accidents, and legal programs [4, 5].

Epidemiological data reveals that individuals suffering from SUD commonly engage in misuse of more than one substance. For instance, in a study, a substantial majority of individuals who met criteria for AUD and OUD - 63.6% and 87.5%, respectively – reported experiencing multiple SUDs [6]. More specifically, alcohol and opioids are often misused together. One study estimates that 33.1% of individuals with OUD also met the criteria for AUD [6, 7]. Co-abuse of opioids and alcohol exacerbates risk, such as increased susceptibility to overdose and individual health hazards, including a higher prevalence of mood and anxiety disorders, as well as intense drug consumption and craving [8]. Thus, understanding the nature of opioid-alcohol misuse is imperative for developing treatment strategies, particularly targeted interventions and public health initiatives, to address unique challenges posed by the co-misuse of opioids and alcohol.

While the co-occurrence of OUD and AUD is well documented, the potential causal relationship between these disorders remains unclear. Opioids primarily activate mu-opioid receptors in the central nervous system, whereas alcohol lacks molecular specificity, interacting with numerous targets throughout the brain [9, 10, 11]. Repeated episodes of heavy alcohol consumption induce central opioid deficiency, mimicking opioid withdrawal [12]. This might sustain alcohol intake through negative reinforcement mechanisms, potentially linking the two disorders [13, 14, 7]. Furthermore, shared genetic factors have been identified among SUDs, suggesting a complex interplay of common and specific biological pathways that influence vulnerability [15]. However, a comprehensive understanding of the common and distinct genetic mechanisms underlying AUD and OUD is lacking.

Here, we integrated public AUD [16] and OUD [17] transcriptomic data from the dorsolateral prefrontal cortex (DLPFC) with genomic data sourced from the latest and most comprehensive GWAS on problematic alcohol use (PAU) [18] and OUD [19] to explore the shared and distinctive molecular basis of these disorders. Our investigation revealed that, in AUD and OUD, there are consistent alterations in several shared biological domains, such as extracellular matrix remodeling, Rho signaling, long coding RNA regulation pathways, and various neuroimmune processes. Despite these parallels, there was minimal convergence at the gene level. Integration of transcriptomic and genomic data reinforced our conclusions and underscored the limited shared heritability of these disorders. This suggests that chronic alcohol and opioid use affect analogous underlying biological mechanisms, albeit distinct genes. This comprehensive approach provides deeper insight into the molecular mechanisms that underpin both disorders, forming a critical foundation for the development of more targeted and effective treatments for individuals affected by them.

## Methods

### Data Collection and RNA-Seq Differential Analysis

Raw fastq files from datasets GSE174409, GSE189139, and PRJNA530758 were obtained using sratoolkit. Details about sample collection, RNA isolation, library preparation, and sequencing can be found in the original manuscripts [16, 17]. Raw sequencing reads underwent adapter trimming and quality control using fastp 57. Samples passing the quality were subjected for the alignment and read count using star to the human genome (GRCh38) with default parameters. Two samples from the OUD dataset (SRR14524862 and SRR14524863) and four from the AUD dataset (SRR16976624, SRR16976667, SRR8843029, SRR8843065) did not pass quality control and were removed from the analysis. Count data representing Alcoholic vs Control condition was organized in matrix format with genes in rows and samples in columns then imported into R version 4.3.1 (https://cran.r-project.org). Counts were within-sample normalized using the counts per million (cpm) function available as part of the “edgeR” package. After, cross-sample normalization was applied using cyclic locally estimated scatterplot smoothing (“loess”) procedure via the “normalizeBetweenArrays” function available as part of the “limma” package. Post normalized values were then scrutinized by Tukey box plot, covariance-based Principal Component Analysis (PCA) scatter plot, and correlation-based clustered heat map to confirm the absence of outliers. For generation of the box plot, PCA scatterplot and the clustered heat map, the “boxplot”, “princomp”, and “pheatmap” functions were used respectively. To remove noise-biased genes, post-normalized values were modeled using locally weighted scatterplot smoothing (lowess) by condition (Coefficient of Variation mean), and the fit plots were inspected. From this inspection, a noise threshold was defined as the lowest mean value at which the linear relationship between mean and Coefficient of Variation is grossly lost. Genes not having at least one sample with a value greater than this threshold were discarded from further analysis with values for surviving genes floored to equal the threshold if less. Filtered and floored values for surviving genes were then tested for differences between conditions using Analysis of Covariance (ANCOVA) under both Akaike Information Criterion (“AIC”) step condition and BenjaminiHochberg (“BH”) False Discovery Rate (“FDR”) Multiple Comparison Correction (“MCC”) procedure. The “Anova” function supported in the “car” package along with commands supported from the “multcomp” and “effect” packages were used to accomplish while the covariates to adjust for were age, pH, gender, post-mortem interval (“PMI”), and RNA integrity number (“RIN”). Testing results were summarized by volcano plot with differential genes identified as those having an ANCOVA BH FDR Corrected P < 0.25 and a linear magnitude difference of means between conditions >= 1.15X. Sample-to-sample relationships were then summarized by covariance-based PCA scatterplot and correlation-based clustered heat map using post noise-filtered and floored values for differential genes only. Corresponding enriched pathways and functions for these genes, along with activation prediction, were obtained using IPA (https://digitalinsights.qiagen.com). Methods described were similarly applied to Opioid count data with the following notable differences: 1) covariates to adjust for were: age, pH, gender, race, PMI, and RIN) differential genes were identified as those having an ANCOVA BH FDR Corrected P < 0.10 and a linear magnitude difference of means between conditions >= 1.25X.

### Rank-rank hypergeometric overlap

RRHO plots were used to compare transcriptomic overlap between OUD1 vs. OUD2, OUD1 vs. AUD, and OUD2 vs. AUD groups. Plots were generated using the RRHO2 package [20], and each gene was scored and ranked by -log10(P-value) multiplied by the sign of the Log2FC. Data visualization was performed using the packages “ggplot2” v3.4.4, “EnhancedVolcano” v1.18.0, “pheatmap” v1.0.12, “eulerr” v7.0.0, and “factoextra” v1.0.7.

### Gene Ontology (GO) analysis

Enrichr was used to query the 2023 GO database of the Gene Ontology Consortium Enrichment criteria were set as *P* -value <0.05 [21].

### Immediate early genes (IEGs) enrichment analysis

Overrepresentation analysis was calculated using the R package phyper with the arguments: **X-1**, the number of DEGs in the IEG gene set; **M**, the total number of genes in the IEG gene set, **N**, the total number of genes absent from the IEG gene set, and **K** is the total number of DEGs. The alpha level was set to 0.05.

### Pathway Enrichment pathway analysis

Canonical pathway analysis was performed using Ingenuity Pathway Analysis (Qiagen, Fredrick, MD) by using each list of DEG as a separate input. Alpha level was set to 0.05 and absolute Z-score >2. Dissimilarity between each pathway was calculated using Jaccard distance and clustered using hierarchical cluster analysis. Dissimilarity matrix was calculated using a custom script while clusters and plots were generated using the R package “factoextra” v1.0.7.

### Upstream regulator analysis

Predicted upstream regulators were identified using Ingenuity Pathway Analysis (Qiagen, Fredrick, MD). Upstream regulators considered for analyses met a corrected P-value < 0.05 and a predicted activation score different from zero. The Z-scores were used to make predictions about the activation status of each modulator.

### Biotype analysis

For gene biotype analysis, DEGs were classified into ten different biotypes (long non-coding RNA, protein-coding, transcribed unitary pseudogene, transcribed unprocessed pseudogene, transcribed processed pseudogene, processed pseudogene, unprocessed pseudogene, miscellaneous RNA, mitochondrial tRNAs, Small nucleolar RNAs) using the R package “biomaRt” v2.56.1 and “org.Hs.eg.db” v3.17.0.

### Partitioned heritability LDSC

Heritability partitioning for OUD and AUD was conducted using the ldsc pipeline [22]. Initially, variants within a 100 Kb interval upstream and downstream of the coordinates of each DEG list, available in the 1000 Genomes European (EUR) phase 3 database, were annotated. Subsequently, chromosome-specific annotation files were generated using the ldsc.py script’s –l2 option, encompassing the gene set within each DEG list. Following this, heritability enrichment was evaluated for the OUD and PAU datasets separately. Moreover, enrichment was also assessed across five other traits: Alcohol Consumption per week and Age of Initiation [23], Major Depressive Disorder [24], Schizophrenia [25] and Neuroticism [26]. An enrichment threshold of 1 and an FDR <0.1 was set, and the enrichment profiles were compared between OUD and AUD datasets.

### Transcriptome-wide association analysis

The MetaXcan software was used to perform a gene-level association study using the AUD and OUD GWAS summary statistics and the psychencode (hg38) transcriptome prediction model (based on 2,188 postmortem frontal and temporal cerebral cortex samples from 1,695 adults) 65. The model used for this association is implemented in the SPrediXcan.py script. In summary, this model assigns weights to individual cis-QTL variants and incorporates measures of variances and covariances of genetic markers in psychencode data to address linkage disequilibrium among the variants [27]. As in the psychencode prediction model the variants were not identified by rs IDs, we included the following tags in the SPredixcan command line –model_db_snp_key varID – keep_non_rsid –additional_output. The predicted gene expression levels significantly associated with the AUD and OUD GWAS were defined based on an FDR 10% threshold. Prior to the gene-level associations study, the markers coordinates were converted from hg19 to hg38 using the liftOver command-line tool from UCSC [28].

## Results

### Differently expressed transcripts in the DLPFC in OUD and AUD

Publicly available RNA sequencing (RNA-seq) data were used to compare the transcriptome-wide changes in the DLPFC of donors diagnosed with OUD [17] and AUD [16]. Phenotypic and demographic information on the analyzed samples can be found in the supplementary material.

Using differential expression analysis with covariate selection, we identified 115 upregulated transcripts (41%) and 81 downregulated (59%; Fig S1A, Table S1) in postmortem DLPFC samples from donors with OUD compared to matched controls. We used hierarchical cluster analysis to group subjects based on differentially expressed genes (DEGs). Control samples formed a single cluster, while OUD samples were divided into two subclusters (Fig S1B). Principal component analysis (PCA) revealed a similar trend (Fig S1C). OUD samples comprised two subgroups: the first, OUD1, was distantly positioned to control samples, while the second, OUD2, was closer to controls. Notably, available phenotypic information proved insufficient to explain OUD heterogeneity. When clustering samples based on the available categorical phenotypic data using Gower distance, they formed two cohesive groups without no clear sub-clustering within groups (Fig S1C). Thus, we conducted a second differential expression analysis using the three identified clusters: control, OUD1, and OUD2. The OUD1 versus control comparison revealed a significantly greater number of DEGs and a larger effect size than the OUD2 versus control comparison (Fig 1A-B; Table S1). Of 196 DEGs identified in the original OUD group, 92% were shared with OUD1, while only 30% were shared with OUD2. Moreover, 62% of DEGs from OUD2 were shared with OUD1, and a subset of 56 genes exhibited differential expression across all group comparisons (Fig 1C). Moreover, 62% of DEGs from OUD2 were shared with OUD1, and a subset of 56 genes exhibited differential expression across all groups (Fig 1C). Correlation analysis among DEGs revealed a moderate correlation between OUD1 and OUD2 groups (R2=0.29, P=2e-16). Additionally, 70 of 72 DEGs common to the OUD1 and OUD2 groups showed changes in the same direction (Fig 1D). Threshold-free correlational analysis using Rank Rank Hypergeometric Overlap (RRHO) confirmed concordant global changes in gene expression between the OUD1 and OUD2 groups (Fig 1E). We further assessed the validity of our groups using hypergeometric analysis of immediate early genes (IEGs). IEGs are activated by drugs of abuse, serving as established markers for neuronal activity in response to psychostimulants [29]. Of 51 analyzed IEGs, 25 were expressed in our samples (Fig 1F, bottom). Although IEGs were enriched in all three clusters, OUD1 exhibited the largest overlapping genes and largest -log10(P-value), followed by the overall OUD group and OUD2 (Figure 1F, top). Together, these findings suggest similar patterns of gene expression among groups, with OUD1 exhibiting heightened statistical power and better representing global changes in gene expression within this OUD population. We used Gene Ontology (GO) and pathway enrichment analysis to understand the differential functional characteristics of each OUD group. OUD1 showed larger number of enriched GO terms than OUD2. Both groups shared 23 enriched GO terms, comprising 7% of OUD1 GO terms and 16% of OUD2 terms (Fig 1G). GO terms common to both groups included functions related to the regulation of miRNA transcription, Tau-protein kinase, and hormone biosynthesis. Molecular functions involved binding of phosphotyrosine residues, transcription regulation, and phosphorylated amino acids (Fig 1G).

**Figure 1:**
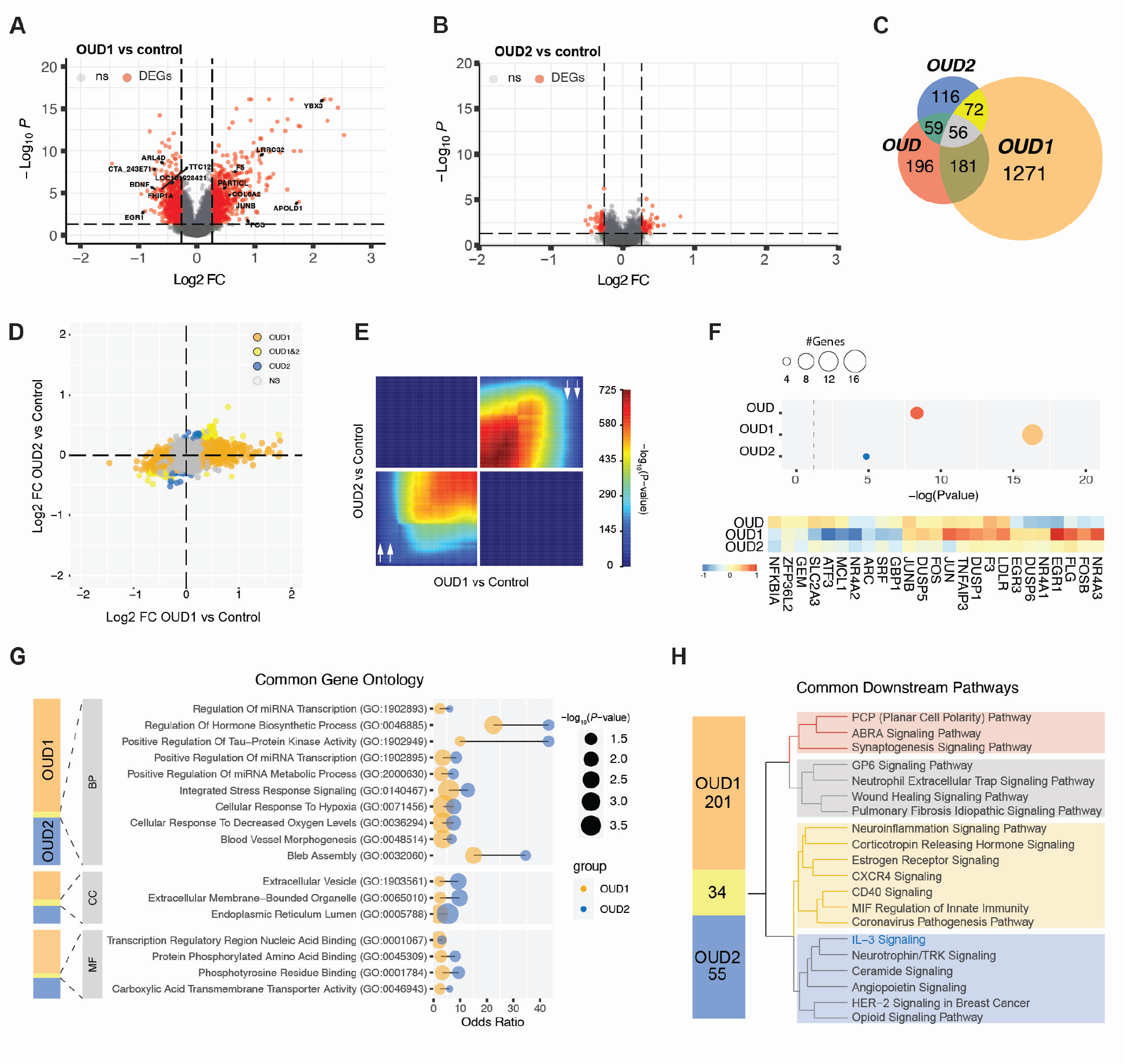
Transcriptomic changes in the DLPFC from donors with OUD. **(A-B)** Volcano plot of RNA-seq data shows DEGs (red) from OUD1 versus unaffected control donors (A) and OUD2 versus control donors (B). The horizontal dashed line represents the FDR significance cut-off (< 0.05), and the dashed vertical lines represent a Log2FC cut-off ± 0.26 (FC=1.2). **(C)** Venn diagram representing the overlap of DEGs in OUD1, OUD2, and OUD groups shows 56 shared DEGs across all group comparisons. **(D)** Correlation analysis of the fold change of combined DEGs from OUD1, (orange) OUD2 (blue). DEGs shared between OUD1 and OUD2 are shown in yellow. **(E)** RRHO analysis shows concordance of gene expression between OUD1 and OUD2 versus control in the DLPFC. Pixels represent overlap between the genome-wide transcriptome of each comparison, with the significance of overlap color-coded. The bottom left quadrant includes co-up-regulated genes, and the top right quadrant includes co-down-regulated genes compared to the control. Bottom right and top left quadrants include oppositely regulated genes. **(F)** Top: Hypergeometric analysis quantifying the enrichment of immediate early genes (IEGs) within the list of DEGs in OUD (red), OUD1 (orange) OUD2 (blue) groups. The X-axis represents –log10(P-value); the size of the circle represents the number of overlapping genes. Bottom: heatmap represents the Log2FC of the 25 IEGs expressed in the DLPFC samples across OUD1, OUD2, and OUD groups. **(G)** Top Gene Ontology (GO) terms shared between OUD1 (orange) and OUD2 (blue) groups. The x-axis represents the odds ratio and measures effect size, and the size of the circle represents the –log10(P-value). **(H)** Top 20 canonical pathways commonly enriched in the OUD1 and OUD2 groups. Pathways were clustered into four distinct groups based on the similarity of overlapping genes.

Exclusive to the OUD1 group were GO terms including “Response to cytokine” (Biological Process), “Cell-cell junction” (Cellular Component), and “CARD domain binding” (Molecular Function). Conversely, unique GO terms for the OUD2 group were “Cyclooxygenase pathway” (Biological Process) and “Secondary Lysosome” (Cellular Component) (Table S2). Overlap between OUD1 and OUD2 became more evident when examined at the pathway level using Ingenuity pathway enrichment analysis.

We found 34 shared pathways, representing 17% of OUD1 and 62% of OUD2 pathways. Shared pathways clustered into four distinct groups based on the number of shared genes, encompassing processes linked to inflammatory response such as Cytokine storm, IL-3, and neuroinflammation, and CXCR4 signaling and to neuronal signaling pathways such as Opioid, Corticotropin-Releasing Hormone, ERK/MAPK, and Synaptogenesis signaling pathways (Fig 1H). Meanwhile, OUD1 showed exclusive enrichment in interleukin pathways including IL-1, IL-6, IL-8, IL-9, IL-10, IL-12, IL-15, and IL-23 signaling. In contrast, pathways exclusive to OUD2 were associated with oxidative stress, including NRF2-mediated oxidative stress response, Fatty acid *β*-oxidation, Oxidative phosphorylation, and Mitochondrial dysfunction signaling pathways (Table S3).

Using a similar approach to compare transcriptome-wide changes within the DLPFC of donors with AUD and well-matched control donors, we identified 159 upregulated (44%) and 205 downregulated (56%; Fig 2A; Table S1) transcripts. Hierarchical cluster analysis revealed two groups. Group 1 captured AUD samples exhibiting large effect sizes, while Group 2 encompassed a mixture of AUD and control samples, lacking clear differentiation (Fig 2B). PCA (Fig 2C) revealed a continuum gradient between the two groups, supporting the cluster analysis. The top enriched GO terms were associated with biological processes of “Cellular Biogenic Amine Catabolic Process,” molecular function “Aromatic Amino Acid Transmembrane Transporter Activity,” and cellular compartment “Collagen-Containing Extracellular Matrix” (Fig 2D; Table S2). One hundred and forty-nine canonical pathways were enriched and clustered into four groups encompassing signaling pathways, including WNT and IL-8 signaling, neuronal pathways such as Serotonin receptors and Long-term potentiation, cellular development pathways like Semaphorin signaling, and finally, the Opioid signaling pathway. (Fig 2E; Table S3).

**Figure 2:**
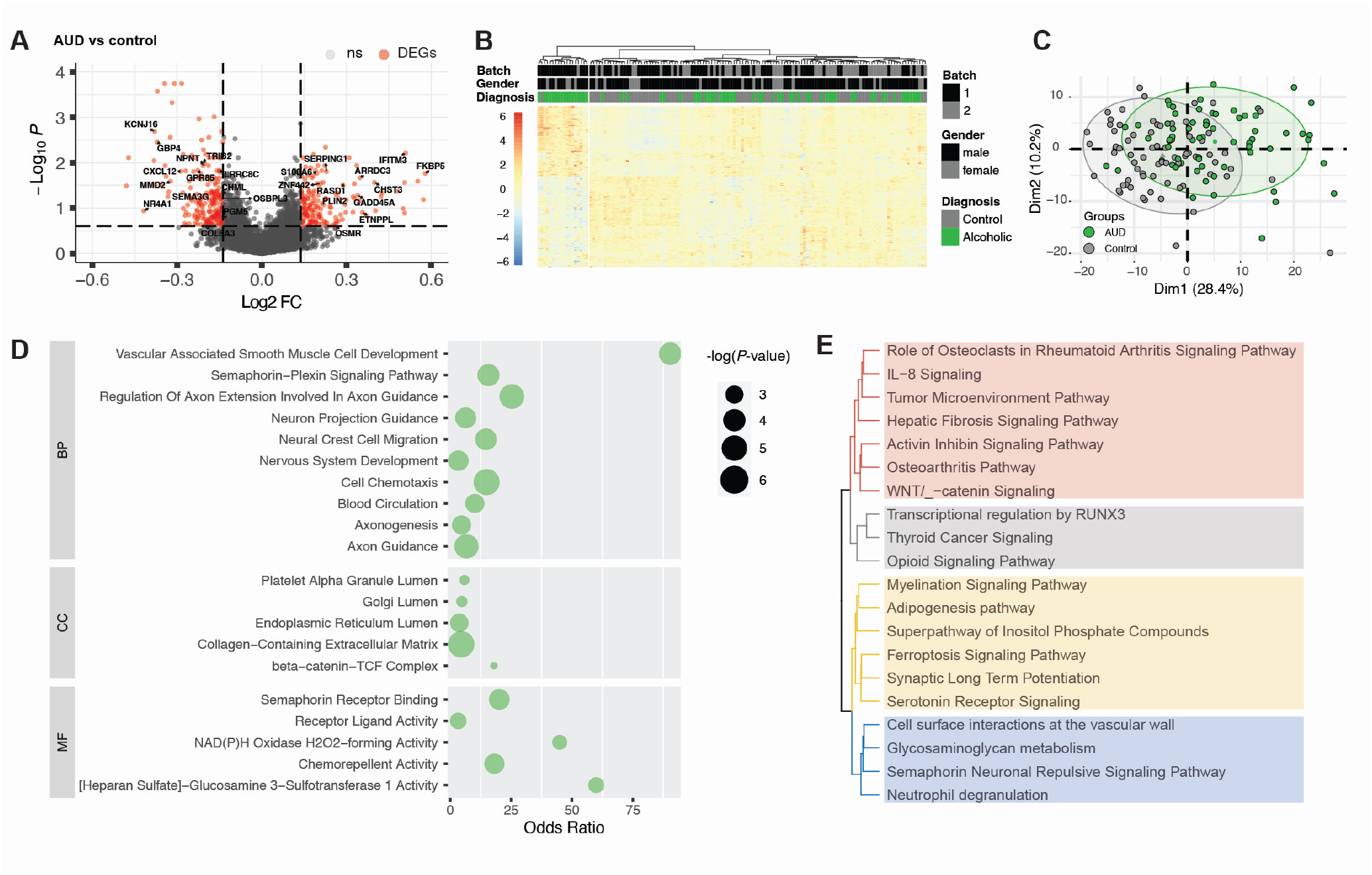
Transcriptomic changes in the DLPFC from donors with AUD. **(A)** Volcano plot shows DEGs (red) from AUD versus control donors in the DLPFC. The horizontal dashed line represents the FDR significance cut-off (< 0.25), and the dashed vertical lines represent a Log2FC cut-off ± 0.20 (FC=1.15). **(B)** Heatmap representing gene expression of all 364 DEGs. Samples were clustered using hierarchical cluster analysis by expression similarity. Each column represents a different subject. Top bar plots represent phenotypic data from each subject. Samples from donors with AUD are shown in green, and the control is in grey. AUD samples are clustered into a single distinct group. **(C)** PCA shows a large overlap between the AUD and control samples. Ellipses represent a 90% confidence range. **(D)** Top GO terms enriched in the AUD group. The size of the circle represents the –log10(P-value), and the odds ratio measures the effect size. **(E)** Pathway analysis shows the top 20 canonical pathways enriched in the AUD group. Pathways were clustered based on the extent of overlap among genes.

### Comparing Transcriptomic changes in the DLPFC of donors with AUD and OUD

We conducted a comparative analysis at both the gene and functional levels of the transcriptomic changes associated with AUD and OUD. The OUD1 group, exhibiting the majority of observed DEGs and a larger effect size, was designated as our primary focus for comparison with AUD samples. Additional comparisons with OUD2 are included in the supplementary material (Table S1-3). RRHO analysis demonstrated significant concordance in the magnitude and direction of the effect size between OUD1 and AUD groups (Fig 3A). OUD2 versus AUD groups also exhibited agreement, albeit with reduced effect size (Fig S2A). Protein-coding genes constituted the majority of DEGs in both AUD and OUD1 groups, with consistent distribution (94% of AUD and 84% of OUD1 DEGs). Long non-coding RNAs (lncRNA) constituted 5% of AUD and 12% of OUD1 DEGs (Figure 3B). Seventy-two DEGs were common to both groups, accounting for 20% of AUD and 6% of OUD1 DEGs. Notably, 92% (66 genes) of the common DEGs exhibited fold changes in the same direction in both groups, with only six genes exhibiting opposite fold changes (Fig 3C-D). Among shared DEGs, 96% were protein-coding genes, and 4% were lncRNAs (*FAM27C, LIFR-AS1*, and *LINC02217*). The predominant categories within protein-coding genes included those that encode enzymes (e.g., *CHML, ETNPPL, FKBP5, RASD1, CHST3, GNA14, GBP4, MTHFD2, ECI2, RAB34, RHOC, ST8SIA6, PGM5*, and *GGT5*), and transporters (*A2M, ABCG2, OSBPL3, S100A6, SLC16A6, SLC38A5, SLC44A3*, and *TAP1*). OUD2 and AUD shared four exclusive DEGs: *TCIM, CHRNA2, EPHB1* (protein-coding genes), and LINC00507 (a non-coding RNA gene). Additionally, six protein-coding genes were shared across all three groups: *MMD2, TTC12, TNS3, KCNJ16, SLC16A6*, and *FHIP1A*. GO analysis revealed DEGs shared between AUD and OUD1 were significantly enriched for terms associated with extracellular matrix, endothelial barrier, chemotaxis, cytokines, and vascular processes (Fig 3E). Complex traits such as SUDs involve hundreds to thousands of genetic loci, each exerting a modest effect [30]. Numerous disease-associated genes contribute to a few common biological processes. To identify enriched associations with specific biological mechanisms, we focus on collective gene effects rather than single gene effects. Ingenuity enrichment analysis of 72 shared genes revealed three clusters of processes: cytokine signaling (e.g., interleukin (IL)-8 and CXCR4 signaling); immune and cancer-associated signaling (e.g. Pulmonary healing and tumor microenvironment pathways); and intracellular signaling pathways (e.g., RHO GTPases) (Fig 3F).

**Figure 3:**
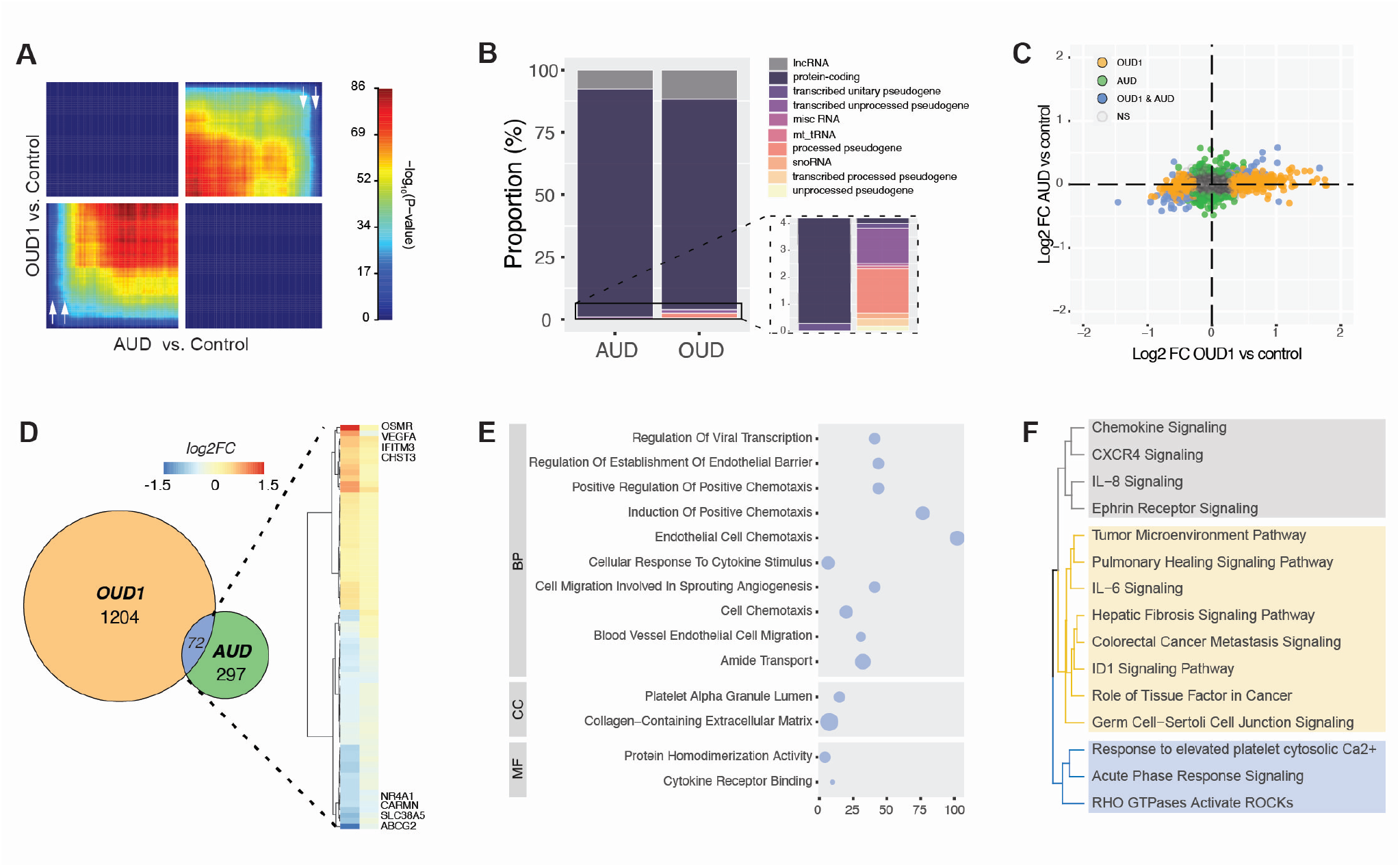
Comparing changes in the DLPFC in donors with AUD and OUD 1. **(A)** RRHO plot shows a large overlap between AUD and OUD1 groups at the genome-wide level. See Fig. 1A for the definition of each quadrant. **(B)** Biotype analysis of all DEGs in the AUD and OUD1 groups reveals the distribution of gene products within each group. The inset provides a closer look at the less common biotypes. **(C)** Scatterplot illustrating the correlation between the fold change of the AUD group (green) and OUD1 group (orange) for all expressed genes in the DLPC. Common DEGs are highlighted in blue, while unchanged changes are represented in gray. **(D)** Venn diagram shows DEGs shared between AUD and OUD1 (blue) along with exclusive DEGs for AUD (green) and OUD1 (orange). The heatmap shows the Log2FC of the 72 shared DEGs between OUD1 and AUD groups. The gene symbols for the top 4 upregulated and downregulated genes in the OUD1 group are highlighted. **(E)** The top GO terms enriched for the common DEGs in both AUD and OUD1 groups are depicted. The circle size reflects the –log10(P-value), and the x-axis illustrates the odds ratio, indicating the effect size. (F) Pathway analysis shows the top 15 canonical pathways enriched for common DEGs in both AUD and OUD1 groups. Pathways were clustered based on the extent of overlap among genes.

### Comparing overlap of canonical pathways enriched in the DLPFC of donors with AUD and OUD

Conducting overrepresentation analysis on overlapping DEGs without considering their effect sizes hinders directional inference. Considering the complex interactions within gene networks and biological processes, changes in a single pathway occur due to alterations in different subsets of genes. Therefore, to evaluate the directionality of changes comprehensively and gain a more nuanced understanding of biological alterations linked to AUD and OUD, we conducted a pathway-level overlap analysis to identify shared and unique pathways in each group. Despite the modest gene-level similarity, there was a notable overlap in biological pathways between the two disorders. Notably, OUD1 and AUD groups shared 51 pathways (34% of OUD1 pathways, 52% of AUD pathways; Fig 4A, left). Among these shared pathways, 31 exhibited a non-zero activation Z-score, enabling us to assess their activation status (Fig 4A, right). Remarkably, among the common pathways, only 4 exhibited a positive activation score in the AUD group, while 30 of 31 pathways showed a positive Z-score in the OUD1 group (Fig. 4A, right). Subsequent correlation analysis categorized pathways into three subgroups based on activation Z-scores. Quadrant 1 contained pathways exhibiting positive Z-scores in both AUD and OUD1 groups. Notably, the Acute Phase Response Signaling Pathway displayed high activation scores (Fig. 4A). Shared DEGs within this pathway were linked to inflammation (*TNFRSF1A, TNFRSF1B, A2M*, and *OSMR*) and the complement system (*C1R* and *SERPING1*) (Fig. 4B). Quadrant 3 showed that only one pathway, RHOGDI Signaling, was suppressed in both groups. Shared DEGs within this pathway included *GNA14, LIMK2*, and RHOCH (Fig. 4B). Quadrant 2 featured pathways activated in the OUD group but suppressed in AUD with varied Z-scores: 1, low absolute Z-scores in both AUD and OUD (e.g., Myelination Signaling); 2, low absolute Z-scores in AUD but high in OUD (e.g., Hepatic Fibrosis Signaling); 3, high absolute Z-scores in AUD but low in OUD (e.g., Phagosome Formation); and 4, high absolute Z-scores in both AUD and OUD (e.g., Pulmonary Healing Signaling). A cluster of pathways showed positive Z-scores in the OUD group and near-zero Z-scores in the AUD group. Among these was the Opioid signaling pathway, reinforcing the recurrent association of opioid neurotransmission dysregulation with both disorders [12, 31]. Further, only 5 pathways were exclusively shared between OUD2 and AUD, whereas 9 pathways were shared across AUD, OUD1, and OUD2 (Table S3). Gene-level analysis confirmed most genes within shared pathways do not overlap between groups despite contributing to dysregulation in the same pathway (Fig. 4B). These results suggest that alcohol and opioids change common biological processes but through dysregulation of distinct gene sets.

**Figure 4:**
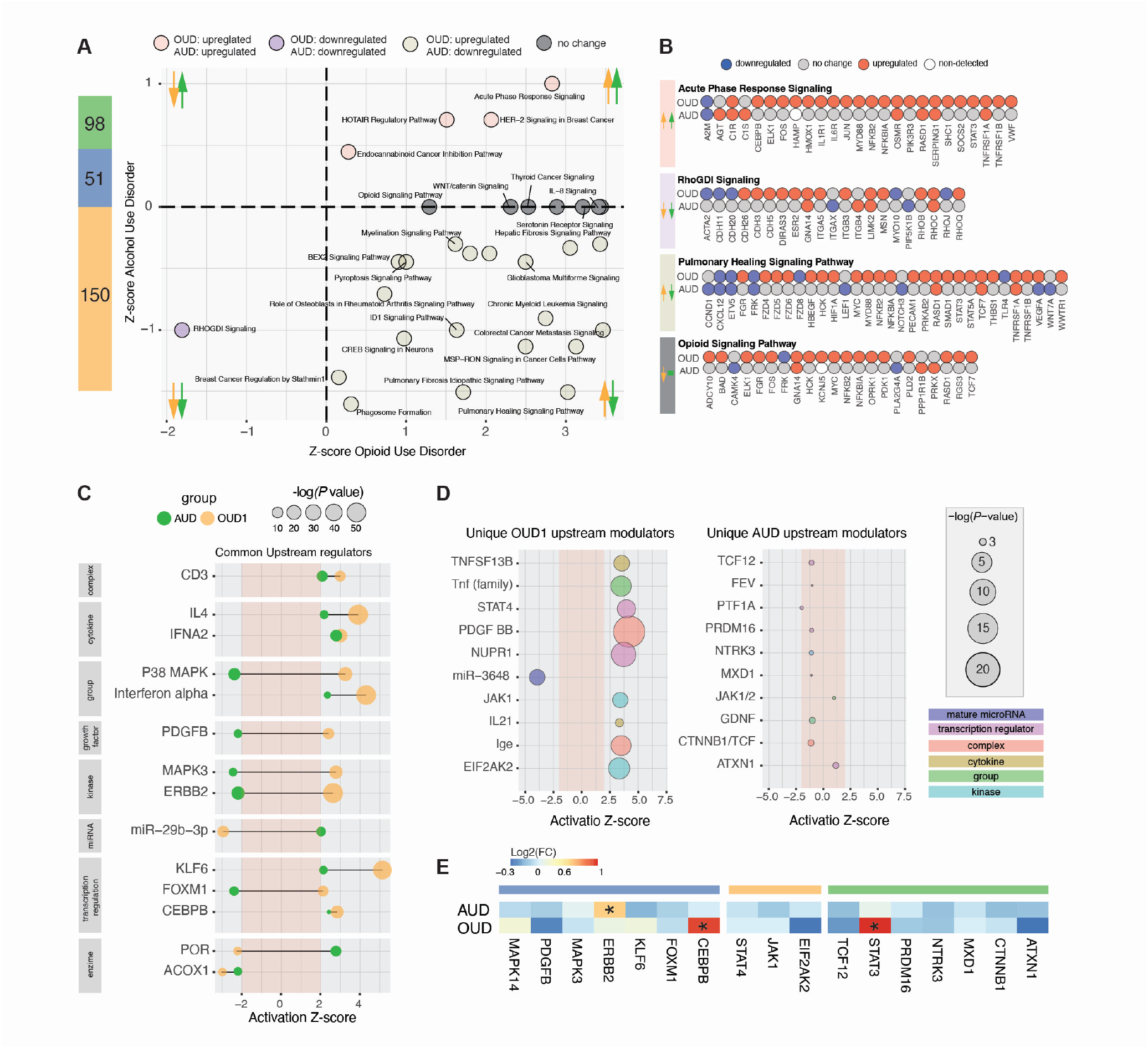
Comparing Transcriptomic changes in the DLPFC of donors with AUD and OUD at the pathway level. **(A)** Left: Bar plot illustrating the percentage of enriched pathways shared between OUD1 (orange) and AUD groups (blue), as well as pathways uniquely enriched in each group. Right: Scatterplot depicting the Z-score correlation of canonical pathways commonly enriched between OUD1 and AUD groups. Pathways were distributed in three quadrants representing their activation status in each group. Pathways with a Z-score > 0 are considered activated, and those with a Z-score < 0 are considered deactivated. **(B)** Gene-level analysis shows down-regulated (blue) and upregulated (red), genes within selected pathways from each quadrant. **(C)** Upstream modulators exhibiting enrichment in both AUD (green) and OUD1 (orange) groups. The x-axis displays the activation Z-score, with the circle size representing the –log10(P-value). Only upstream regulators with an absolute Z-score greater than 2 are included in the plot. The red-shaded area denotes regions with non-significant Z-scores. **(D)** Upstream modulators enriched exclusively in OUD1 (on the left), or AUD (on the right) groups are depicted. The x-axis indicates the activation Z-score, and the circle size corresponds to the –log10(P-value). The red-shaded area shows zones with non-significant Z-scores. Notably, none of the upstream modulators exclusive to AUD reached Z-score significance. **(E)** Heatmap representing the Log2FC of protein-coding upstream modulators within the AUD and OUD1 groups. Cells marked with an asterisk indicate differentially expressed genes.

We also examined pathways uniquely enriched in each disorder. Among those exclusive to the AUD group, 69% displayed a non-zero Z-score, with 65% showing decreased activation. The Galpha (i) signaling events pathway displayed the most significant negative Z-score, while the Dopamine Degradation Pathway exhibited the highest positive Z-score (Fig S2B). Conversely, pathways exclusively enriched in the OUD1 group all showed increased activation, with 65% displaying a non-zero activation Z-score. Pathways linked to inflammatory response (iNOS, IL-9, and IL-23 signaling) and cell differentiation (FGF and EGF signaling) exhibited the most significant positive activation Z-scores (Fig. S2B). Within the 16 pathways exclusively enriched in the OUD2 group, two exhibited a non-zero Z-score: NRF2-mediated Oxidative Stress Response and Mitochondrial Dysfunction (Table S3). A gene-level analysis revealed that DEGs from pathways exclusive to one group were underrepresented in the other. For instance, OUD1 lacked DEGs associated with the AUD-exclusive pathways, like Dopamine Degradation and G alpha (i) signaling events, while AUD lacked DEGs from OUD1-exclusive pathways, such as iNOS and IL-9 signaling (Fig S2C).

### AUD and OUD share upstream modulators associated with inflammatory processes

Gene expression is regulated by upstream mediators including transcription factors and synthetic or natural compounds. Upstream mediators target many interconnected genes or biological pathways, providing valuable insights into gene networks and holding significant therapeutic potential. We employed upstream enrichment analysis to identify potential modulators of DEGs associated with AUD and OUD. Fourteen upstream modulators were commonly enriched in both disorders (P ≤ 0.01 and absolute Z Score ≥ 2). Following pathway-level trends, we found that most of the shared upstream modulators exhibited opposing changes between AUD and OUD1 groups (Fig 4C). Of these, most belonged to the class of transcriptional regulators (21%), followed by group of molecules, kinases, cytokines, and enzymes (14% each). Two mediators of transcriptional response previously implicated in drug addiction were identified: MAPK3 (Mitogen-Activated Protein Kinase 3) and p38 MAPK (p38 mitogen-activated protein kinases) [32, 33]. Interestingly, inflammation modulators like IL4 and interferon alpha showed significant positive Z-scores in both groups, corroborating findings from pathway analyses and previous literature (Fig 4C). Half of the shared upstream modulators showed changes in the same direction in both groups, indicating potential targets for common therapeutical approaches.

Several upstream regulators were exclusively enriched in only one group (Fig 4D). In the OUD1 group, significant enrichments with notable effect sizes were found in modulators primarily associated with inflammation processes, including STAT4, JAK1, IL-21, and members of the Tnf family (Fig 4D). All upstream modulators exclusive to OUD exhibited upregulation, except for microRNA miR-3648, which was predicted to undergo significant inhibition. Conversely, none of the modulators exclusive to AUD exhibited a significant activation Z-score. Modulators associated with AUD, albeit with borderline scores, were linked to inflammation, such as STAT3 and JAK1/2, and to cellular differentiation and proliferation, including TCF12 (transcription factor 12), CTNNB1 (catenin beta 1), and GDNF (Glial cell line-derived neurotrophic factor) (Fig 4D).

We then assessed the expression status of the genes encoding exclusive and shared upstream modulators within AUD and OUD1 groups. Among the 26 enriched upstream modulators classified as gene products, 19 (73%) were expressed in our DLPFC samples (Fig 4E), underscoring the robustness of our findings. Notably, within this subset, STAT3 and CEBPB displayed exclusive differential expression in OUD1, while ERBB2 exhibited exclusive differential expression in AUD. STAT3 and CEBPB were upregulated in both groups, consistent with upstream analysis (upregulated in both groups), whereas ERBB2 showed a change in the opposite direction (Fig 4E; Table S4). Additionally, four signaling processes – P38MAPK, MAPK3, ERBB, and PDGF were consistently upregulated in OUD1 in both upstream (Fig S2B) and pathway analyses (Fig 4A). OUD2 and AUD shared two exclusive upstream modulators— Sirtuin 6 (SIRT6) and norepinephrine — exhibiting a significant negative Z-score in both groups. Five upstream modulators were shared with the OUD1 group, including TNF and miR-3648. CEBPB was the sole upstream regulator shared across all three groups, showing an increased Z-score in OUD1 and AUD and a negative Z-score in OUD2 (Table S4).

### Multiomics analysis reveals limited genomic and transcriptomic overlap between AUD and OUD

Gene expression alterations can stem from environmental adaptations or inherent genetic predispositions. To identify causal relationships and genes associated with risk factors for these disorders, we integrated transcriptomic data using two methodologies. First, we used Linkage disequilibrium score regression (LDSC) to estimate the proportion of the heritability attributable to our subset of DEGs using summary statistics from GWAS encompassing PAU, OUD, and other neuropsychiatric disorders. We found a significant contribution of the AUD-DEG list to the heritability explained by the PAU and DrinksPerWeek GWAS summary statistics. Similarly, we observed a significant association between genes identified from the OUD1-DEGs list and the OUD GWAS (Fig 5A). While there was no enrichment for OUD1 genes in the AgeOfInitiation (alcohol) GWAS, it is noteworthy that the trend in the P-value was consistent (P=0.11; Fig 5A). However, neither AUD nor OUD1 showed significant enrichment for Major Depressive Disorder (MDD), Schizophrenia, and Neuroticism (Fig 5A).

**Figure 5:**
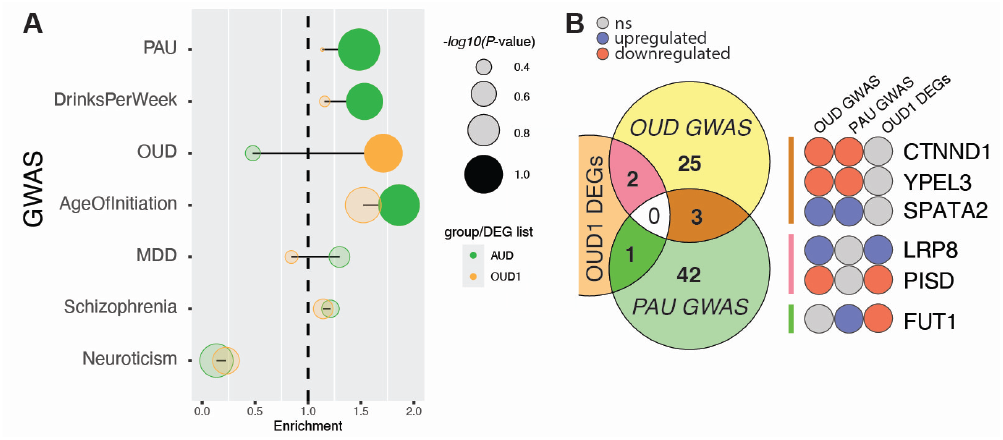
Genomic and transcriptomic integration. **(A)** Extent of heritability elucidated by AUD and OUD1-DEGs across various traits and neuropsychiatric conditions. **(B)** Transcriptome-wide association analysis showing predicted gene expression levels within the DLPFC based on genetic variants associated with PAU and OUD and their overlapping with the set of OUD1-DEGs. Displayed are the genes shared across multiple conditions, accompanied by their predicted and observed gene expression levels. Genes showing upregulation are highlighted in red, while those exhibiting downregulation are depicted in blue.

Our second approach used Transcriptome-wide association studies (TWAS) to predict gene expression levels within the DLPFC based on PAU and OUD genetic variants. Forty-six genes significantly linked to PAU and 30 to OUD were found. Additionally, three genes(*CTNND1, YPEL3*, and *SPATA2*)showed significant associations with both GWAS (Fig 5B). Two genes enriched in the OUD *GWAS, LRP8* and *PISD*, exhibited differential expression in the OUD1 group, while one gene linked to the PAU GWAS, *FUT1*, showed differential expression in the OUD1 group. Remarkably, LRP8 and PISD exhibited consistent expression patterns across analyses (Fig 5B). Based on predicted expression levels from OUD and PAU GWAS, *CTNND1*, and *YPEL3* showed upregulation, while *SPATA2* showed downregulation in both GWAS datasets. No overlap was found between AUD-DEGs and TWAS-enriched genes.

## Discussion

Despite studies examining the molecular basis of AUD and OUD, there remains a limited understanding of the unique and shared molecular mechanisms underlying both disorders. Our study comparing the transcriptome changes within the DLPFC of donors with OUD and AUD revealed distinct alternations in individual genes but convergence on specific processes and biological pathways. Within shared biological pathways, numerous genes downregulated in AUD exhibit no change in OUD, while other genes were upregulated in OUD but exhibited no change in AUD. Thus, though the influence of these drugs on the transcriptome converges to some extent on a few common biological processes, the mechanism of these changes is likely specific to each drug and reflects unique molecular signatures underlying AUD and OUD. The similarities observed at both pathway and functional levels could hold significance for the high prevalence of co-occurring AUD and OUD, potentially contributing to comparable behavioral phenotypes within substance use disorders (SUDs) or elucidating the physiological ramifications of chronic substance use.

DEGs common to both groups were mostly protein-coding genes, followed by lncRNAs, and associated with various circulatory and vascular processes, chemotaxis, and endothelial functions. The Collagen-Containing Extracellular Matrix (GO:0062023) emerged as the GO process with the most significant P-value, ranking highest in significance within the AUD group and second within the OUD1 group but not reaching significance in OUD2. All shared genes within this GO domain exhibited changes in the same direction in the AUD and OUD groups. Research suggests that ECM remodeling directly influences neuroadaptations associated with chronic opioid exposure, while perineuronal nets, specialized ECM structures surrounding inhibitory neurons, may contribute to compulsive drinking behaviors in AUD through the mediation of reward memories [34]. These findings underscore the intricate relationship between ECM remodeling and addiction, illustrating the adaptable interplay between these systems in response to addictive drugs.

Pathway-level analysis revealed divergent regulation among specific pathways and convergence on others. The Acute Phase Response signaling (APR) and HOTAIR regulatory pathways exhibited co-upregulation in the AUD and OUD1 groups. RHOGDI signaling pathway exhibited co-downregulation. Pathways exhibiting concordant alterations in both groups present potential therapeutic targets for both disorders.

APR signaling is critical in mediating inflammation and is implicated in SUD pathophysiology [35]. The metabolism of addictive substances, including opioids and alcohol, produces reactive oxidative species (ROS), inducing an inflammatory response and triggering cytokine production in peripheral circulation [36]. Peripheral inflammatory signals activate neuroimmune signaling, potentially contributing to SUD development through broad impacts on synaptic organization, neurotransmission, neuronal networks, and behavior [37, 38].

LncRNAs are increasingly recognized as essential links between clinical phenotypes and GWAS findings [39]. HOTAIR, an important regulatory RNA, regulates the inflammatory response via NF-kb modulation. In a recent study, HOTAIR induction in immune cells was required for the pro-inflammatory response [40]. Upregulated HOTAIR expression is involved in CNS disorder pathogenesis by sponging miRNA, inducing apoptosis, activating microglia and neuroinflammation, and inhibiting BDNF transcription [39]. These findings reinforce evidence that neuroinflammation plays a crucial role in the development and persistence of SUDs.

The RHOGDI signaling pathway, a negative regulator of RHO GTPases, was the sole pathway downregulated in both groups. RHO GTPases modulate diverse processes, including cell proliferation, apoptosis, cytoskeletal reorganization, and membrane trafficking via actin regulation. Actin remodeling plays a role in synaptic formation, function, and plasticity, while changes to synaptic plasticity may underlie the neurobiology of addiction [41]. RHOGDI signaling likely regulates morphological and neurochemical plasticity associated with substance use. Reversing changes to synaptic plasticity associated with addiction may serve as a novel treatment for treating SUDs [42, 43].

The opioid signaling pathway was significantly enriched in both groups. However, while it showed a significative positive Z-score in the OUD1 group, it displayed a zero Z-score in the AUD group. This divergence stems from the fact that although there was a significant overrepresentation of DEGs associated with opioid signaling in AUD, there was no clear bias toward up or downregulation. Given naltrexone, an opioid receptor antagonist, is a first-line treatment for AUD, it is unsurprising that endogenous opioid signaling was associated with AUD [44, 45]. However, clinical studies on naltrexone efficacy have variable results. In one study, 20-55% of patients prescribed naltrexone experienced relapse, with another 16-60% discontinuing use [46, 47]. Within the opioid signaling pathway, GNA14 (G protein subunit alpha 14) was the sole shared gene between AUD and OUD groups, showing downregulation. GNA14 encodes a member of the alpha q subfamily of G proteins and initiates a cascade that activates protein kinase C (PKC) through beta-type phospholipase C (PLC-*β*) enzymes. Studies in mice indicate genetic deletion of PLC-*β* or pharmacological inhibition of Gq enhances morphine’s antinociceptive effects [48]. Our findings suggest a synergistic approach to modulating the endogenous opioid system may hold promising treatment options for patients with AUD and OUD.

Numerous pathways demonstrated unique transcriptomic alterations within OUD1 and AUD groups. In AUD, pathways associated with monoamine degradation (i.e., Dopamine, Noradrenaline, Adrenaline, and Serotonin) were predominantly upregulated. Monoamine neurotransmitters are crucial in regulating alcohol intake and reward. Enzymatic metabolism, mediated by enzymes like Monoamine Oxidase A (MAOA) and Catechol-O-methyltransferase (COMT), modulates monoamine levels [49]. Dopamine can also be metabolized by MAOB and Aldehyde Dehydrogenase. Notably, genes encoding these enzymes were exclusively upregulated in AUD, suggesting a hypomonoaminergic state in the DLPFC of donors with AUD. The dopamine hypothesis suggests multiple repeated drug use induces homeostatic changes in the brain, leading to a hypodopaminergic state, a hallmark of SUDs [50]. Evidence from clinical and preclinical studies supports reduced dopaminergic activity in brain areas like the NAc and PFC following chronic drug use [51, 52, 53]. Differences in the dopamine states between AUD and OUD groups may stem from variations in substance use recency. For instance, in this study, the cause of death of individuals within the AUD group was linked to repercussions of chronic alcohol consumption rather than acute alcohol intoxication. Conversely, in the OUD group, all deaths were attributed to overdose incidents. Consequently, toxicological factors could impact dopamine dynamics and gene expression, contributing to observed differences.

OUD1-exclusive pathways showed enrichment for diverse inflammatory processes, consistent with literature demonstrating heightened immune activity in the brains of individuals with OUD. Studies have shown although chronic alcohol use generally elicits a pro-inflammatory response, the pro or anti-inflammatory cytokine profile may vary in response to different stages of AUD progression and recovery [54]. Importantly, the OUD1 group exhibited a significantly larger effect size than the AUD one. Consequently, outcomes of the non-overlapping pathway analysis should be interpreted cautiously due to insufficient statistical power. Notably, OUD-1 specific pathways not only differed from those observed in the AUD group but also exhibited distinctions compared to the OUD2 group. This subgroup consists of donors diagnosed with OUD, exhibiting gene expression patterns characterized by smaller effect sizes than OUD1. These findings suggest that individuals in the OUD2 group may be either in an earlier stage of the disease than those in the OUD1 group or might possess a compensatory mechanism at the gene expression level to maintain brain homeostasis in response to chronic opioid exposure. These insights are significant in clinical contexts and warrant careful consideration in future analyses, especially with the prevalence of studies utilizing large datasets.

Half of the upstream modulators enriched in both AUD and OUD exhibited consistent alterations. These modulators were related to immune and transcriptional functions, with half exhibiting consistent alterations. Transcription factors have the potential to govern extensive gene networks as such, may be candidates for therapeutic intervention. Notably, transcription factors KLF6 (Kruppel-like factor-6) and CEBPB (CCAAT Enhancer Binding Protein Beta), which play pivotal roles in modulating neurodegeneration and inflammation, were upregulated in AUD and OUD1. KLF6 (Kruppel-like factor 6) facilitates axon extension and myelination by promoting pro-myelinating signals, thereby regulating the development of oligodendrocytes from oligodendrocyte progenitor cells [55, 56]. On the other hand, CEBPB (CCAAT Enhancer Binding Protein Beta) is involved in regulating pro-inflammatory genes in microglia and has been implicated in Alzheimer’s disease progression [57, 58]. Thus, while KLF6 activation might foster axon regeneration in response to inflammation and neurodegeneration induced by chronic drug exposure, CEBPB activation may serve as a master regulator of neuroinflammation, alongside IL4 and interferon alpha, both of which are commonly upregulated in both AUD and OUD groups.

The robustness of our findings is accentuated through multilevel independent analysis, revealing several crucial biological components that consistently surfaced across genes, pathways, and upstream levels. Such a comprehensive approach holds promise for devising more efficacious treatments for OUD and AUD. For instance, common upstream modulators like CEBPB or KLF6 could be considered potential therapeutic targets. By targeting these modulators, it may be possible to manipulate specific gene networks associated with biological pathways such as HOTAIR and APR signaling, which play pivotal roles in various pathologies of AUD and OUD, including neuroinflammation.

These robust correlations highlight the fundamental importance of these biological processes in shaping the respective disorders, thus reinforcing the credibility and coherence of our study’s findings. Furthermore, the absence of a robust shared heritability across disorders serves to underscore these results. While genes differentially expressed in AUD demonstrated a significant capacity to elucidate the heritability of PAU, Age Of Initiation, and Drinks Per Week, those associated with OUD1 accounted for a substantial portion of the heritability specific to OUD. Consequently, despite overarching similarities at the pathway level, our findings unveil distinct mechanisms within transcriptional networks governing alcohol and opioid dependencies, ultimately giving rise to diverse neuroadaptations.

Some limitations should be considered in the interpretation of the findings in this study. The transcriptomic data are based on two independent public RNA-seq datasets from separate studies. Consequently, the samples originated from different sources and underwent distinct processing methods. Moreover, the sequencing procedures employed diverse platforms and sequencing parameters. Therefore, the potential impact of variances in brain bank sources, processing techniques, and sequencing platforms should be acknowledged. The correlational nature of the comparison analysis is an inherent limitation, given the different data sources, making a direct comparison challenging. Thus, the findings should be considered within the context of these limitations, and further studies with standardized procedures across multiple datasets may be warranted to validate and extend the current observations. To address these challenges, we made rigorous efforts to minimize potential confounding factors. Specifically, each dataset included matched pairs of control samples, and both datasets were subjected to the same quality control standards and analysis pipeline to enhance consistency and comparability. The strength and validity of our findings were also reaffirmed by the consistent results observed upon integrating these data with independent genomic datasets.

Overall, these findings suggest misuse of alcohol or opioids affects different mechanisms, as seen in differences in single-gene transcriptional alterations, but ultimately, these disparate mechanistic alterations converge onto similar biological pathways. Our results align with the current literature implicating inflammation and immune function dysregulation from alcohol and opioid misuse. Moreover, just as this study aimed to expand upon and explore the current understanding of the genetic underpinnings of SUD by utilizing transcriptomics, epigenomics offers a way to examine factors influencing alterations in the transcriptome. Research has begun to examine candidate genes of SUD that have been identified in GWAS for potential post-translational modifications [59, 60, 61, 62]. By investigating epigenetic mechanisms, such as methylation, acetylation, histone modifications, and chromosome remodeling, to name a few, we may be able to identify molecular mechanisms that drive changes in gene transcription and lead to aberrant neuroadaptations and circuitry seen in SUD.

## Acknowledgments

This work was supported in part through the computational and data resources and staff expertise provided by Scientific Computing and Data at the Icahn School of Medicine at Mount Sinai, supported by the Clinical and Translational Science Awards (CTSA) grant UL1TR004419 from the National Center for Advancing Translational Sciences, and by the grant K01AA028292 from NIH to RLK. PASF is the beneficiary of a Maria Zambrano Grant of the University of Leon funded by the Ministry of Universities (Madrid, Spain) and financed by the European Union-Next Generation EU. Support also was provided by a Merit Award (I01 BX004820) from the Department of Veterans Affairs.

---

**Figure 6:**
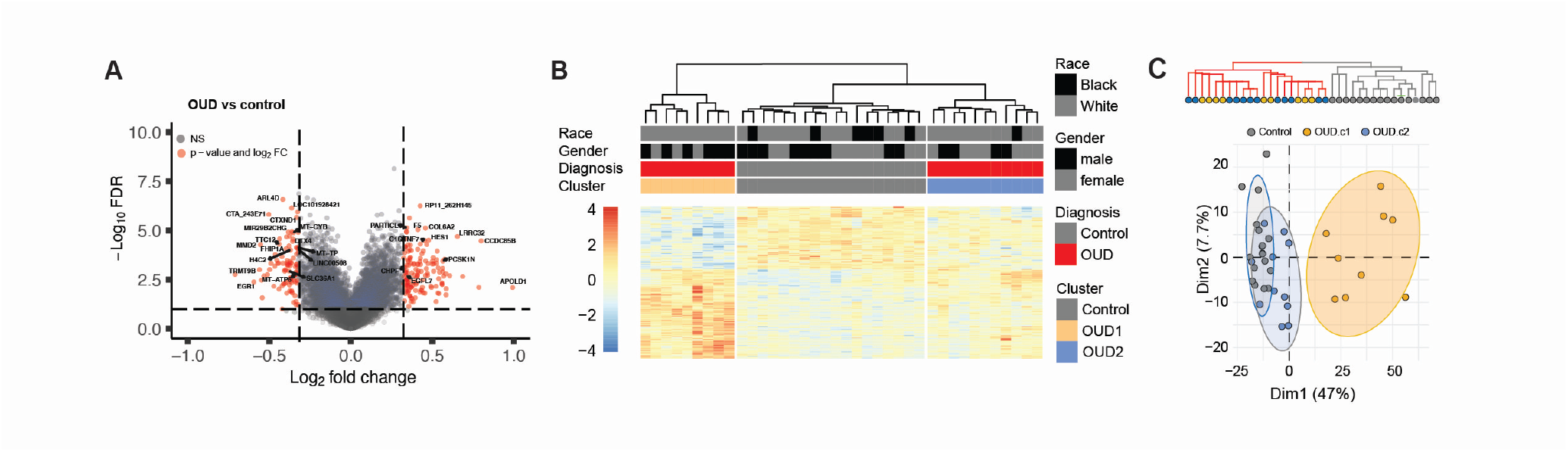
Heterogeneity in the gene expression pattern within OUD samples. **(A)** Volcano plot shows DEGs (red) from OUD versus healthy control donors in the DLPFC. The horizontal dashed line represents the FDR significance cut-off (< 0.05), and the dashed vertical lines represent a Log2FC cut-off ± 0.26 (FC=1.2). **(B)** Heatmap representing gene expression of all DEGs. Samples were clustered using hierarchical cluster analysis by expression similarity. Each column represents a different subject. The top bar plots represent phenotypic data from each subject. OUD donors are shown in red, and control in grey. OUD samples clustered into two distinct groups, OUD1 (orange) and OUD2 (blue). **(C)** Top: Cluster analysis using categorical phenotypic data shows the existence of two distinct clusters, control (gray) and OUD (red). Bottom: Principal component analysis showing the separation of the OUD samples in reduced space. Ellipses represent a 90% confidence range for OUD1 (orange), OUD2 (blue), and Control (grey) groups.

**Figure 7:**
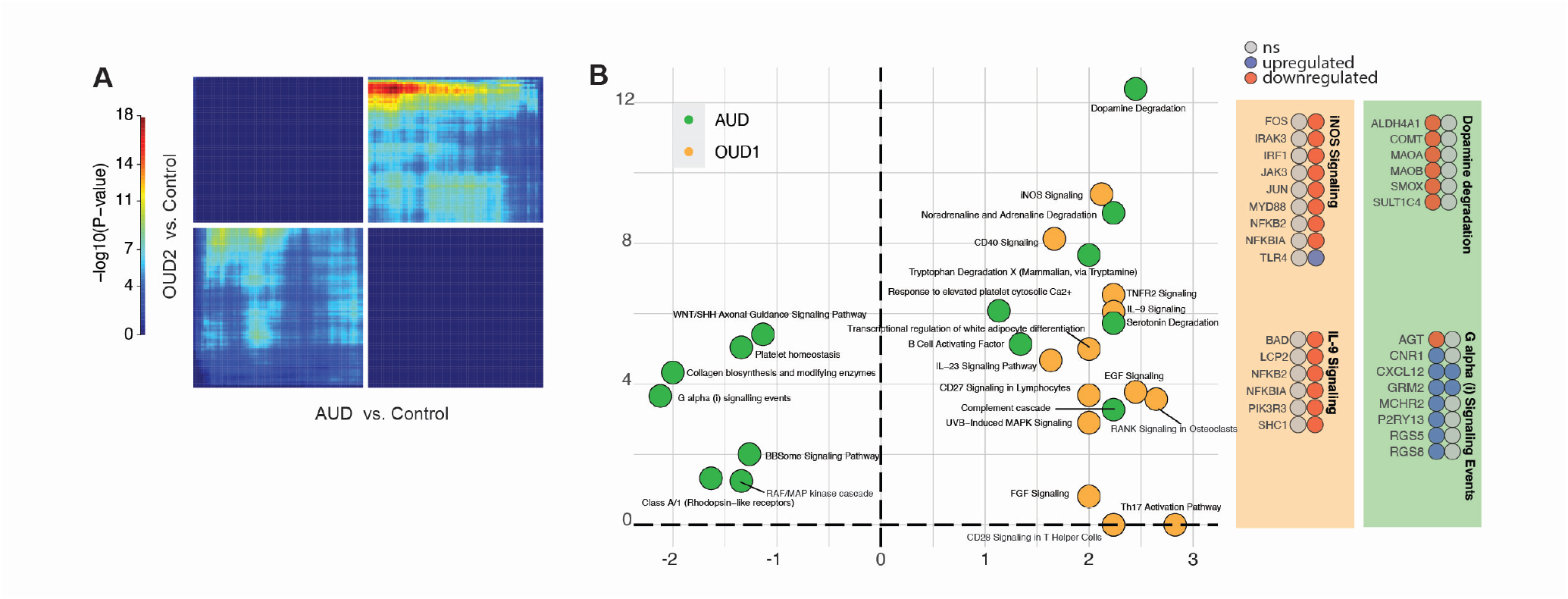
Transcription-wide and pathway-level comparisons between AUD and OUD groups. **(A)** RRHO analysis shows limited concordance of gene expression between OUD2 and AUD versus control in the DLPFC. Pixels represent overlap between the genome-wide transcriptome of each comparison, with the significance of overlap color-coded. The bottom left quadrant includes co-up-regulated genes, and the top right quadrant includes co-down-regulated genes compared to the control. The bottom right and top left quadrants include oppositely regulated genes. (**B, left)** Scatterplot illustrating pathways exclusively enriched in OUD1 (orange) and AUD (green) groups. The x-axis displays the activation Z-score, while the y-axis shows -log10(P-value). Only pathways with a non-zero Z-score are included in the plot. **Right**, Gene-level analysis displays down-regulated genes (blue), upregulated genes(red), non-changed genes (gray), and undetected genes (white) within selected pathways.

